# Unusual actin-binding mechanism and the role of profilin in actin dynamics of trypanosomatid parasites

**DOI:** 10.1101/2023.01.06.522972

**Authors:** Andrea Vizcaíno-Castillo, Tommi Kotila, Konstantin Kogan, Ryuji Yanase, Juna Como, Lina Antenucci, Alphee Michelot, Jack D. Sunter, Pekka Lappalainen

## Abstract

Diseases caused by *Leishmania*, and *Trypanosoma* parasites, such as leishmaniasis and African sleeping sickness, are a major health problem in tropical countries. Due to their complex life cycle involving both vertebrate and insect hosts, and > 1 billion years of evolutionarily distance, the cell biology of these trypanosomatid parasites exhibits pronounced differences to animal cells. For example, the actin cytoskeleton of trypanosomatids is highly divergent when compared to the other eukaryotes. To understand how actin dynamics are regulated in trypanosomatid parasites, we focused on a central actin-binding protein profilin. Co-crystal structure of *Leishmania major* actin in complex with *L. major* profilin revealed that, although the overall folds of actin and profilin are conserved in eukaryotes, *Leishmania* profilin contains a unique α-helical insertion, which interacts with the target binding cleft of actin monomer. This insertion is conserved across the Trypanosomatidae family, and is strikingly similar to the structure of WH2 domain, a small actin-binding motif found in many other cytoskeletal regulators. We demonstrate that the WH2-like motif contributes to actin monomer-binding and enhances the actin nucleotide exchange activity of *Leishmania* profilin. Surprisingly, unlike other profilins characterized so far, *Leishmania* profilin inhibited formin-catalyzed actin filament assembly in a mechanism that is dependent on the presence of the WH2-like motif. By generating profilin knockout and knockin *Leishmania mexicana* strains, we show that profilin is important for efficient endocytic sorting in parasites, and that the ability to bind actin monomers and proline-rich proteins, as well as the presence of a functional WH2-like motif, are important for the *in vivo* function of *Leishmania* profilin. Collectively, this study uncovers the molecular principles by which actin dynamics are regulated by profilin in trypanosomatids. Moreover, the unusual actin-binding mechanism of profilin identified here could be applied for designing inhibitors against pathogenic trypanosomatid parasites.

**AUTHOR SUMMARY:** *Leishmania* and *Trypanosoma* parasites are a major health problem as they cause various diseases in humans and other vertebrates. Currently, there are no specific drugs to treat the diseases caused by these trypanosomatid parasites. Similar to all other eukaryotes, trypanosomatid parasites have an actin cytoskeleton, which is essential for the viability of parasites. Interestingly, both actin and actin-regulatory machineries of these parasites are highly divergent from the ones of animals, making them possible drug targets to treat diseases caused by these parasites. To uncover how the actin cytoskeleton of trypanosomatid parasites is regulated, we focused on a central actin-binding protein, profilin. Importantly, our experiments revealed that trypanosomatid profilins interact with actin through a different structural mechanism as compared to animal profilins, and have specific effects on the assembly of actin filaments. Our genetic studies demonstrate that these specific features of trypanosomatid profilin are also critical for the proper function on this protein in parasites. Our study provides new insight into the cell biology of trypanosomatid parasites. We also envision that the structural and functional differences between trypanosomatid and human profilins can be applied for developing compounds for selectively neutralizing *Leishmania* and *Trypanosoma* parasites.

## INTRODUCTION

The eukaryotic *Leishmania* parasites are the etiological agents of a group of diseases collectively known as leishmaniases. These parasites cause three main types of infections: cutaneous leishmaniasis, visceral leishmaniasis, also known as kala-azar, and mucocutaneous leishmaniasis. The severity of disease varies from asymptomatic to fatal if untreated. Despite a significant amount of research conducted on these parasites, leishmaniasis remains an important health problem with approximately 700,000-1,000,000 new cases per year [1, 2]. The related *Trypanosoma* parasites also cause severe diseases, such as African sleeping sickness and Chagas disease. Both *Leishmania* and *Trypanosoma* genera belong to the Trypanosomatidae family of flagellated protozoan parasites [3].

*Leishmania* parasites have a complex life cycle that involves both insect vectors (sand flies) and mammalian hosts with differentiation into two major cell morphologies: flagellated, motile promastigotes and macrophage-resident, non-motile amastigotes [4]. Due to their unusual life cycle, and because the family of trypanosomatid parasites diverged early in the evolution from other eukaryotes [5], they exhibit biological peculiarities that deviate them from those of well-studied organisms, such as animals, yeasts, and plants. An interesting example of such biological features is their atypical cytoskeleton, which is considered to be mainly microtubule-based [6]. Also the actin is present in trypanosomatid parasites, albeit its role seems to be limited to a subset of cellular processes [7-9]. In animals, the actin cytoskeleton contributes to several cellular functions including motility, morphogenesis, adhesion, vesicular traffic, cytokinesis, and endocytosis [10, 11], but according to current understanding, actin and actin-regulating proteins in trypanosomatids are mainly involved in endocytosis, vesicular trafficking, and assembly of flagellum, and hence important for the viability of these parasites [12-14]. In *Leishmania*, actin may additionally be involved in kinetoplast remodeling during cell division [15].

Actin is an abundant protein, which is conserved throughout evolution from Asgard archaea to all eukaryotes [16]. Globular actin monomers (G-actin) can spontaneously assemble into helical filaments (F-actin) in which actin subunits arrange in a head-to-tail orientation, creating two structurally distinct filament ends known as the barbed end and the pointed end. Furthermore, actin molecules bind an adenosine nucleotide, either ATP or ADP. Actin filament assembly occurs predominantly through incorporation of ATP-actin monomers to the filament barbed end, and filament disassembly occurs mainly through dissociation of ADP-actin monomers from the filament pointed end. Filament turnover is powered by ATP, because in the filamentous form, actin catalyzes ATP hydrolysis, whereas ADP in the actin monomer can be exchanged for ATP to ‘recharge’ the monomer for a new round of filament assembly. The coordinated actin filament polymerization produces pushing forces that, for example, promote formation of plasma membrane protrusions for cell migration and plasma membrane invaginations during endocytosis. In order to control actin filament assembly and disassembly in space and time, a large repertoire of actin-binding proteins (ABPs) evolved to regulate different aspects of actin dynamics. These include the Arp2/3 complex, which catalyzes nucleation of branched actin filament networks for endocytosis and cell migration, formins, which assemble linear actin filaments for other cellular processes, as well as a large array of proteins controlling actin filament disassembly and cytoplasmic actin monomer pool [11, 17].

*Leishmania* and *Trypanosoma* actins show ∼70% sequence identity to vertebrate actins, which makes them among the most divergent actins in the eukaryotic lineage [14]. Despite the large divergence in sequence, a recent structural and biochemical study on *L. major* actin (LmActin) demonstrated that the conformation of actin filaments in *Leishmania* is nearly identical to their vertebrate counterparts. However, due to differences in the subunit-subunit interfaces, the parasite actin filaments display more rapid turnover compared to animal actin filaments. The same study also revealed that *L. major* cofilin fragments LmActin filaments more frequently compared to mammalian cofilin, demonstrating that both actin filaments and their interplay with cofilin display pronounced differences between animals and trypanosomatids [18].

Along with cofilin, a handful of other canonical ABPs are present in *Leishmania* and *Trypanosoma* species [14, 19]. These include the small actin monomer-binding protein profilin, which is found in all eukaryotic organisms that contain a regulated actin cytoskeleton. In animals, yeasts and plants, profilins inhibit spontaneous nucleation of actin filaments, promote elongation of pre-existing actin filament barbed ends, and prevent assembly of actin monomers to the filament pointed ends [11]. Profilins can also accelerate the ADP-to-ATP nucleotide exchange on actin monomers [20-22]. Besides binding actin, profilins also interact with poly-proline rich motifs, which are typical for actin filament nucleating/polymerizing proteins such as formins [23]. Formins are composed of formin homology 1 (FH1) and formin homology 2 (FH2) domains, which consists of poly-proline stretches and interact with actin to promote filament assembly, respectively. Formins contain also other domains involved in regulating their subcellular localization and activity [24]. Interaction of profilin with the formin FH1 domain allows the delivery of actin monomer/profilin complexes to the adjacent FH2 domains. Hence, upon activation, formins typically work in synergy with profilin to accelerate actin filament assembly in cells [25-27]. Profilin expression was also demonstrated in trypanosomatids [28-30], but the mechanism by which this protein controls actin dynamics in *Leishmania* and *Trypanosoma* parasites has remained largely unclear. An earlier study reported that *L. donovani* profilin binds poly-proline peptides and accelerates nucleotide exchange on rabbit muscle actin. Moreover, depletion of profilin was reported to affect cellular growth and endocytic trafficking [30], and contribute to mitotic spindle orientation and cell cycle progression in *L. donovani* parasites [31]. However, no structural information of trypanosomatid profilins is available, and all biochemical work so far has been performed by using a heterologous combination of mammalian and *Leishmania* actin-binding proteins, which were recently demonstrated to be unfavorable substrates for each other [18]. Thus, the mechanisms by which trypanosomatid profilins interact with actin monomers, and regulate actin dynamics together with other proteins, such as formins, have remained elusive.

To uncover how trypanosomatid profilins control actin dynamics, we determined the crystal structure of *L. major* profilin (LmProfilin) in complex with LmActin. Although the overall folds of actin and profilin are conserved in evolution, our structural work revealed that *Leishmania* profilin harbors a peculiar WH2 domain –like α-helix, which makes contact with actin. Biochemical and genetic studies revealed that this insertion, which is conserved across the Trypanosomatidae family, is important for high-affinity actin monomer binding and nucleotide exchange *in vitro*, as well as for the proper function of profilin in endocytosis in *Leishmania* parasites. Moreover, we provide evidence that, unlike profilins from other eukaryotes, *Leishmania* profilin inhibits formin-catalyzed actin filament assembly through a mechanism that is dependent on the WH2-like motif. These findings demonstrate that the actin monomer – profilin interplay is divergent in trypanosomatid parasites as compared to animals, and propose that the specific structural features of actin-profilin interactions may serve as good targets for selectively neutralizing *Leishmania* and *Trypanosoma* parasites.

## RESULTS

### Crystal structure of *Leishmania* profilin-actin monomer complex

To elucidate the mechanism by which trypanosomatid profilins interact with actin monomers to control cytoskeletal dynamics, we expressed and purified recombinant LmProfilin and LmActin, and studied their interactions by X-ray crystallography. We obtained crystals of the *Leishmania* profilin in complex with an ATP-actin monomer, and determined the structure of the complex at 2.2 Å resolution. The structure revealed that *Leishmania* profilin forms a 1:1 stoichiometric complex with ATP-actin monomer and interacts with the barbed end face of the actin monomer at the border of actin subdomains 1 and 3 similar to other profilins (Fig. 1A). The overall fold of *Leishmania* profilin is similar to the determined structures of profilins from other organisms, such as mammals, yeasts, malaria parasite, and Asgard archaea [21, 32-36]. Furthermore, the conformation of *Leishmania* actin in the complex with profilin is very similar to the structure of actin in the previously determined from bovine and Archaea profilin-actin complexes (Fig. S1A). Interestingly, the actin-binding interface of profilin is not particularly well-conserved between the *Leishmania* and mammalian proteins, apart from certain key residues discussed later in the text (Fig. S1B).

**Figure 1.**
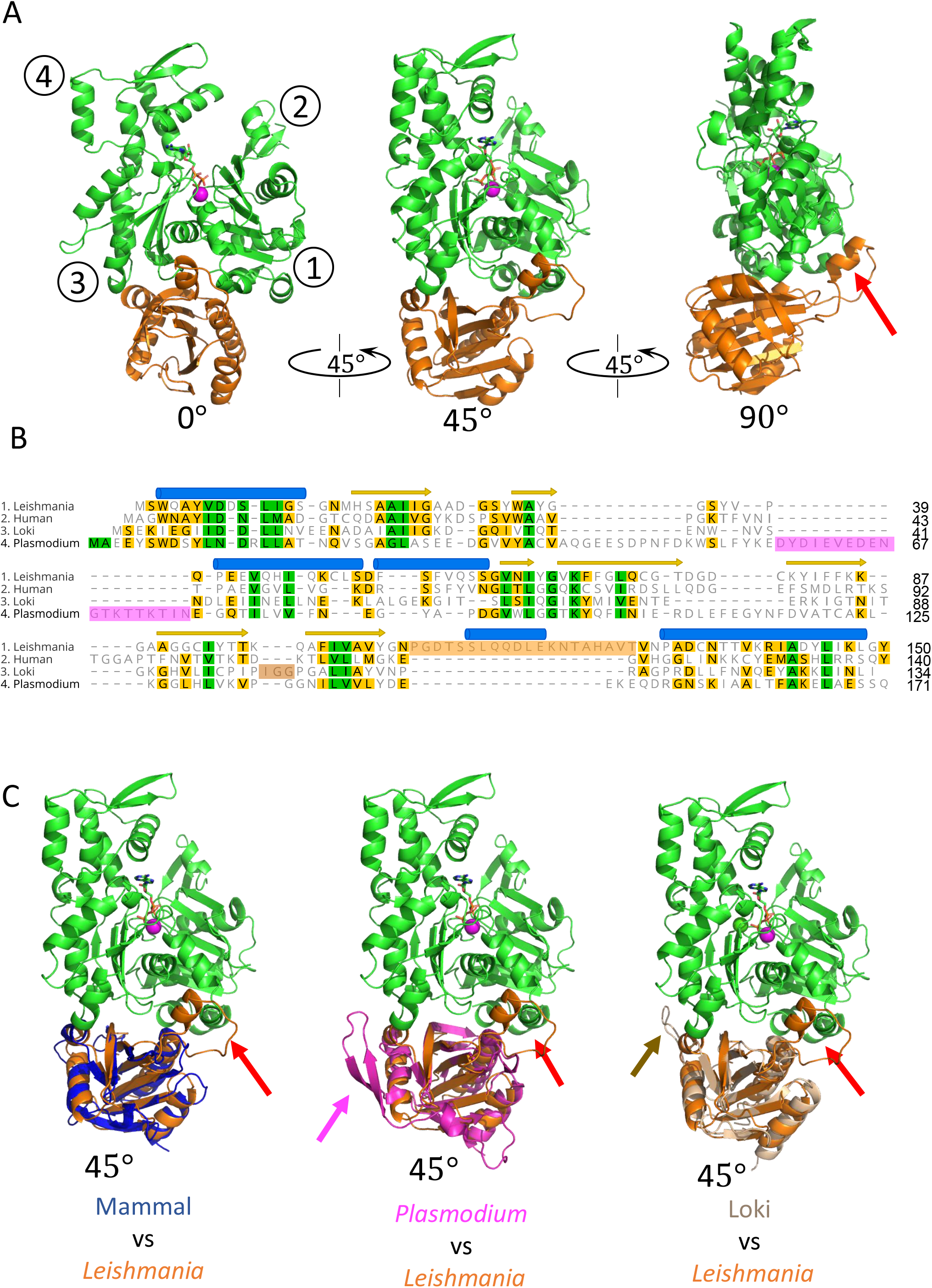
Crystal structure of *L. major* profilin in complex with actin. **(A)** Three orientations (0°, 45°, 90°) of the co-crystal structure of LmProfilin (orange) - actin (green) complex. The ATP nucleotide (brown) and the associated Ca2+ ion (magenta ball-shaped) in actin are highlighted. The sub-domains of actin are labeled by 1-4 numbers in circles. The specific insertion in *Leishmania* profilin is indicated with a red arrow in the panel on right. **(B)** Structure-based protein sequence alignment (performed by Dali server [68]), of profilins from trypanosomatid parasite (*L. major*, PDBID: 8C47, chain B; UniProt: Q4Q5N1), human (*Homo sapiens*, PDBID: 6NBW, chain C; UniProt: P07737; [69]) Asgard archaea (Lokiarchaeum, PDBID: 5ZZB, chain B; UniProt: A0A0F8V8L2; [36]), and malaria parasite (*Plasmodium falciparum*, PDBID: 2JKG, chain A; UniProt: Q8I2J4; [70]). Specific insertions, which are not present in mammalian profilin, are highlighted by pink in *Plasmodium*, by orange in *Leishmania*, and by light brown in Loki profilin sequences. **(C)** Superimposition of the profilins from human, *Plasmodium* and Asgard archaea with the *Leishmania* profilin, when in complex with an actin monomer. The positions of *Leishmania*, malaria parasite and Lokiarchea –specific insertions in the structures are indicated with red, pink and light brown arrows, respectively. A view of 45° was selected to better visualize the locations of the insertions in all three profilins.

The most striking difference of *Leishmania* and *Trypanosoma* profilins compared to the profilins from other organisms is the presence of a ∼20 amino acids insertion [28, 29]. In our crystal structure, this insertion is located between the last β-strand and C-terminal α-helix of the profilin fold and it adopts an α-helical conformation flanked by small stretches of flexible linker sequence. Interestingly, the α-helical part of the loop interacts with the target binding cleft of actin, between actin subunits 1 and 3. (Fig. 1A, B). The loop insertion appears to be specific for trypanosomatid profilins. This is because although also malaria parasite and Loki archaea profilins contain short insertions as compared to mammalian profilins, those are located at different positions of the profilin fold, and interact with different surfaces of an actin monomer (Fig. 1B, C). The specific loop in *Plasmodium* profilin interacts mainly with the barbed end side of subdomain 3 surface, whereas the Loki-loop targets the same subdomain, but at the back surface.

Thus, the crystal structure of *Leishmania* profilin-actin complex shows that trypanosomatid profilins interact with actin monomers through a mechanism that is distinct from those of profilins from other organisms.

### Trypanosomatid profilins harbor a WH2-like actin-binding motif

More detailed analysis of the binding mode of trypanosomatid-specific loop of profilin with actin shows that two leucines, Leu115 and Leu119, of the α-helical region of the insertion associate with the hydrophobic pocket of actin formed by residues Ile345, Leu346, Ile349 and Tyr143 in the target binding cleft located between subdomains 1 and 3 (Fig. 2A, B). Interestingly, the position of *Leishmania* profilin α-helix on the actin monomer, and the mechanism by which it interacts with the surface of actin, are very similar to that of the WASP homology-2 (WH2) domain (Fig. 2C). WH2 domain is a short, ubiquitous motif of 15-20 amino acids present in many regulators of actin filament dynamics. These include, for example, actin filament nucleation promoting factors WASP, N-WASP and WAVE complex, as well as proteins catalyzing actin filament nucleation/polymerization, such as Leiomodin, Spire, Cobl and Ena/VASP [37]. Typically, WH2 domains utilize two conserved leucines or isoleucines in their α-helical region for interactions with actin, and these hydrophobic residues are also present in the α-helical region of the loop insertion of *Leishmania* profilin (Fig. 2D). In most WH2 domains, the α-helix is followed by another region called LKKV or LRRV motif (Leu-Lys/Arg-Lys/Arg-Val), which, however, is absent from *Leishmania* profilin. Interestingly, also other regulators of actin that interact with the barbed end surface apply similar mechanistic properties in their mode of actin-binding. For example, the ADF-H fold of cofilin, gelsolin and twinfilin contain an α-helix, which inserts in a similar fashion to the target binding cleft between actin subdomains 1 and 3. However, in contrast to the canonical WH2 domains and the α-helical WH2 domain –like insertion of *Leishmania* profilin, the α-helices in these proteins have opposite orientation. In cofilin, twinfilin and gelsolin domains, the N-terminus of the α-helix is facing towards the pointed end of actin, whereas in the WH2 domains and in the α-helical insertion of *Leishmania* profilin, the N-terminus of α-helix is facing towards the barbed end of actin (Fig. 2C). Thus, different regulators of actin have found similar ways to interact with actin monomers through convergent evolution.

**Figure 2.**
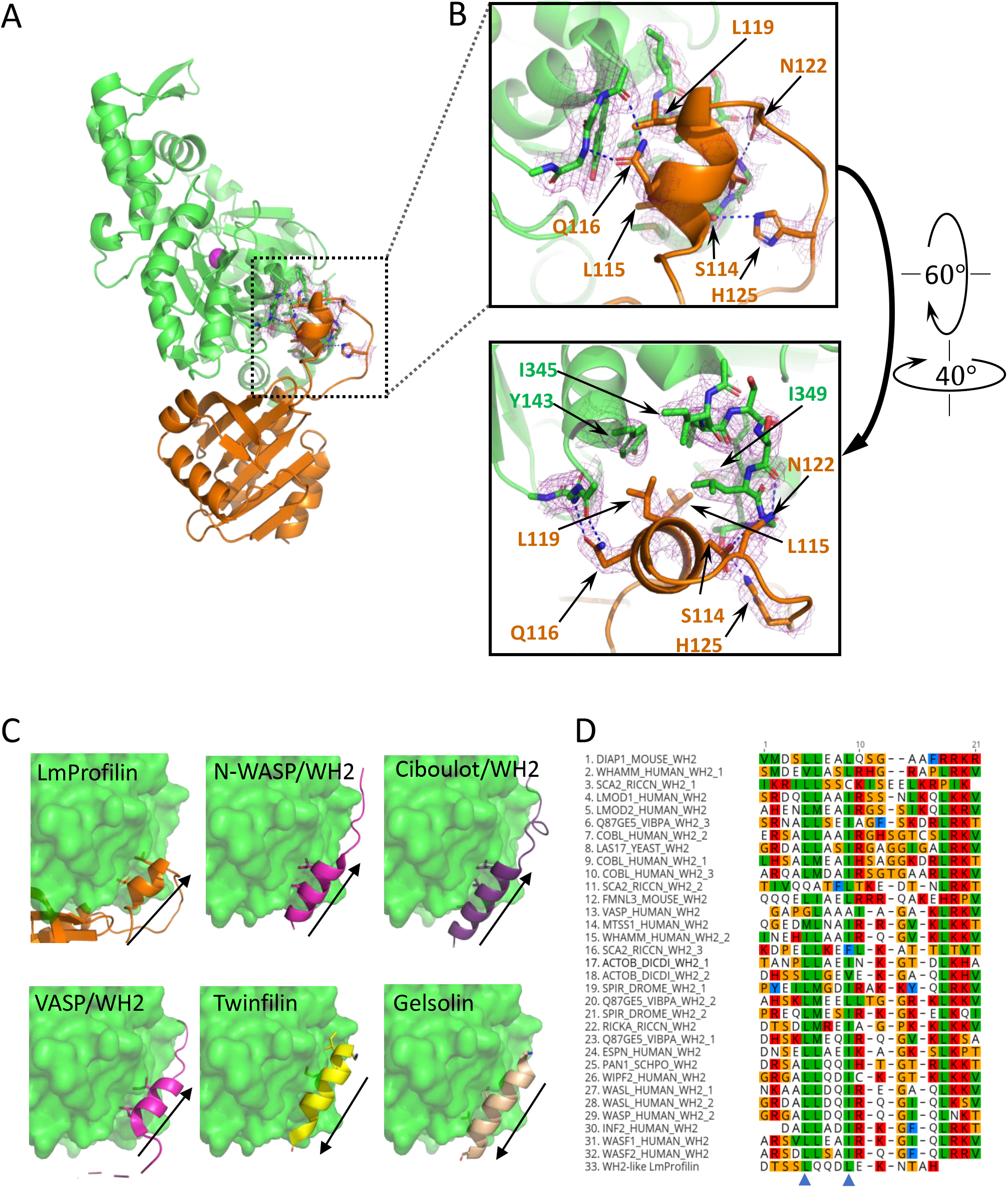
Trypanosomatid parasites harbor a WH2 domain –like α-helical insertion. **(A)** Side view of the *Leishmania* profilin-actin complex showing the interactions of profilin α-helical insertion with actin. **(B)** Magnified view of the interactions of *Leishmania* profilin α-helix with actin. Key residues contributing to the interaction are marked by sticks and labeled with the same color as the corresponding protein molecule. Electron density map (2*F*_0_ – *F*_C_, *σ* = 1.0) is also shown for selected residues. **(C)** 737 Comparison of α-helical regions of *Leishmania* profilin insertion (PDBID: 8C47), selected WH2 domains [N-WASP/WH2 (PDBID:2VCP; [71]), Ciboulot/WH2 (PDBID:1SQK; [72]), and VASP/WH2 (PDBID:2PBD; [73])], as well as twinfilin (PDBID:3DAW; [74]), and gelsolin (PDBID:1T44; [75]). The positions of key hydrophobic residues involved in actin-binding are shown. Arrows indicate the direction of polypeptide chain from the N-terminus to the C-terminus of α−helix. **(D)** Multiple sequence alignment of selected WH2 domains and the α-helical insertion of *Leishmania* profilin. The critical hydrophobic residues mediating actin monomer binding in WH2 domains are indicated with blue arrowheads.

We next examined the role of the WH2 domain–like structural motif (hereafter termed the ‘WH2-like motif’) in actin monomer binding by LmProfilin. For these experiments, we generated five mutant versions of LmProfilin and examined their binding to *L. major* ATP-actin monomers by isothermal titration calorimetry (ITC). In the mutant versions of profilin the entire WH2-like motif was deleted (LmProfilin-ΔWH2), or the two conserved leucines of WH2-like motif in contact with actin were replaced by serines (LmProfilin-WH2-SS). Moreover, we introduced two mutations (LmProfilin-K68A and LmProfilin-K86E) to the ‘main’ actin-binding interface, which is conserved in all profilins. The equivalents of these two mutations in other organisms were reported to affect actin monomer binding to different extents [38]. Finally, we introduced a mutation to the putative poly-proline binding site of the protein (LmProfilin-Y6A), which is not in contact with actin monomer (Fig. 3A). ITC experiments showed that the interaction between LmActin with LmProfilin produced an exothermic reaction and the binding isotherms were best fit to one-site binding model. Wild-type LmProfilin and LmProfilin-Y6A mutant bound ATP-actin monomers with high affinity (K_d_∼80 nM Fig. 3B, C; Fig. S2), whereas mutations at the main binding interface either completely abolished actin-binding (LmProfilin-K86E), or resulted in a very low affinity binding to LmActin (LmProfilin-K68A; K_d_ ∼ 6 *μ*M) (Fig. 3C; Fig. S2). Interestingly, the mutant proteins in which the key actin-interacting residues of the WH2 motif were substituted by serines, or harbored complete deletion of the motif, still bound actin monomers, although with ∼ 15-20 fold reduced affinity as compared to the wild-type profilin (Fig. 3B,C and Fig. S2).

**Figure 3.**
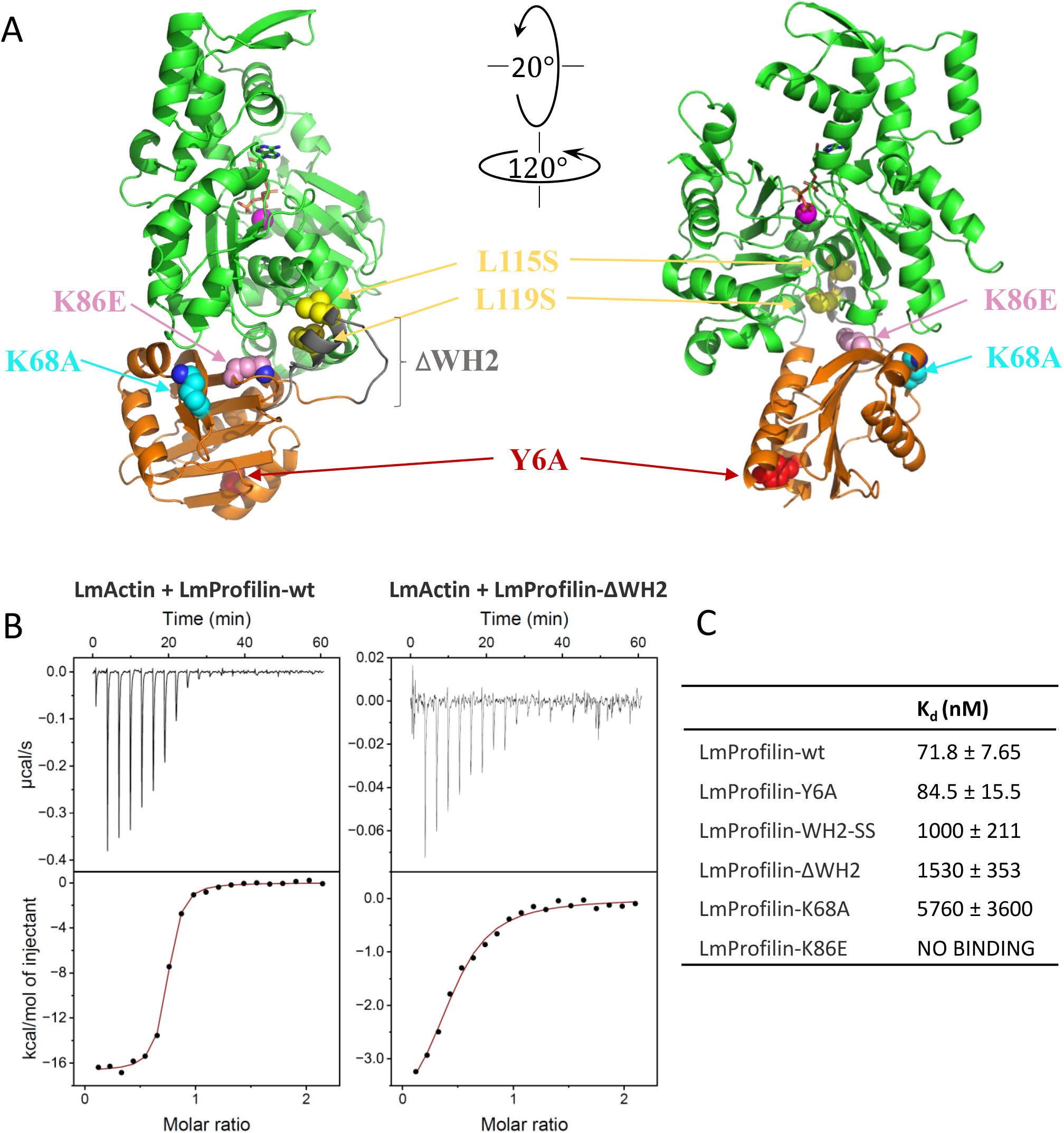
Site-directed mutagenesis reveals the roles of different protein motifs of *Leishmania* profilin in actin-binding. **(A)** The location of amino acid residues that were mutated in *Leishmania* profilin (yellow, L115 and L119; red, Y6A; gray, deleted WH2 motif; pink, K86; cyan, K68) are indicated in the profilin-actin complex (shown in two different orientations). **(B)** Examples of the data from the isothermal titration calorimetry assay. Baseline corrected thermograms (upper graphs) and integrated data fit to one-site binding model (lower graphs) are shown. **(C)** Dissociation constants (K_d_, in nM) of wild-type and mutant *Leishmania* profilins from *Leishmania* ATP-actin monomers, obtained by ITC experiments.

Together, the structural and mutagenesis data provide evidence that LmProfilin harbors a WH2–like motif, which interacts with the hydrophobic cleft between actin subdomains 1 and 3. The WH2-like motif is not essential for interaction of *Leishmania* profilin with monomeric actin, but increases the affinity of profilin for actin. Importantly, this insertion and its key actin-binding residues are conserved in *Leishmania* and *Trypanosoma* parasites, as well as in other trypanosomatids (Fig. S3), indicating that profilins from all trypanosomatid species apply the unique WH2-like motif to regulate actin dynamics.

### *Leishmania* profilin binds proline-rich proteins and catalyzes nucleotide exchange on actin

Most profilins catalyze nucleotide exchange on actin monomers, and a recent study provided evidence that *L. donovani* profilin can accelerate nucleotide exchange on rabbit muscle actin to some extent [30]. To examine the possible effects or LmProfilin on the rate of nucleotide exchange of LmActin monomers, we monitored the ATP-ATTO-488 fluorescence anisotropy kinetics in the presence of actin and wild-type/mutant profilins using the approach described in [39]. Unlike rabbit muscle actin, LmActin monomers do not efficiently exchange their bound nucleotide for ATP-ATTO-488 (Fig. 4A). The presence of *Leishmania* profilin lifts this inhibition and promotes rapid nucleotide exchange in a dose-dependent manner. This effect is observed for about 400-700 seconds, after which anisotropy signals slowly decrease instead of reaching steady state values (Fig. 4B). Since injecting an additional dose of ATP-ATTO-488 after the anisotropy signals have returned to low values does not result in the formation of a new peak, we interpret this effect as a possible gradual inactivation or unfolding of nucleotide-free monomeric Mg-actin during the exchange reaction (Fig. S4A) [40, 41]. Please note that in the experimental conditions of nucleotide exchange assay, the concentrations of ADP and ATP-488-ATTO are very low and there is no unlabeled ATP. Consistent with its lower affinity for actin, the ΔWH2 profilin mutant comparatively showed reduced nucleotide exchange efficiency. The K68A mutant has no exchange efficiency, demonstrating that the conserved actin-binding interface of profilin is essential for nucleotide exchange in the *Leishmania* protein (Fig. 4C). Interestingly, the interplay between actin and profilin seems to have co-evolved, as yeast (*Saccharomyces cerevisiae*) profilin showed no efficiency in promoting *Leishmania* actin nucleotide exchange (Fig. S4B); conversely, the catalytic activity on rabbit actin monomers decreased progressively with the degree of divergence of the profilin (Fig. S4C).

**Figure 4.**
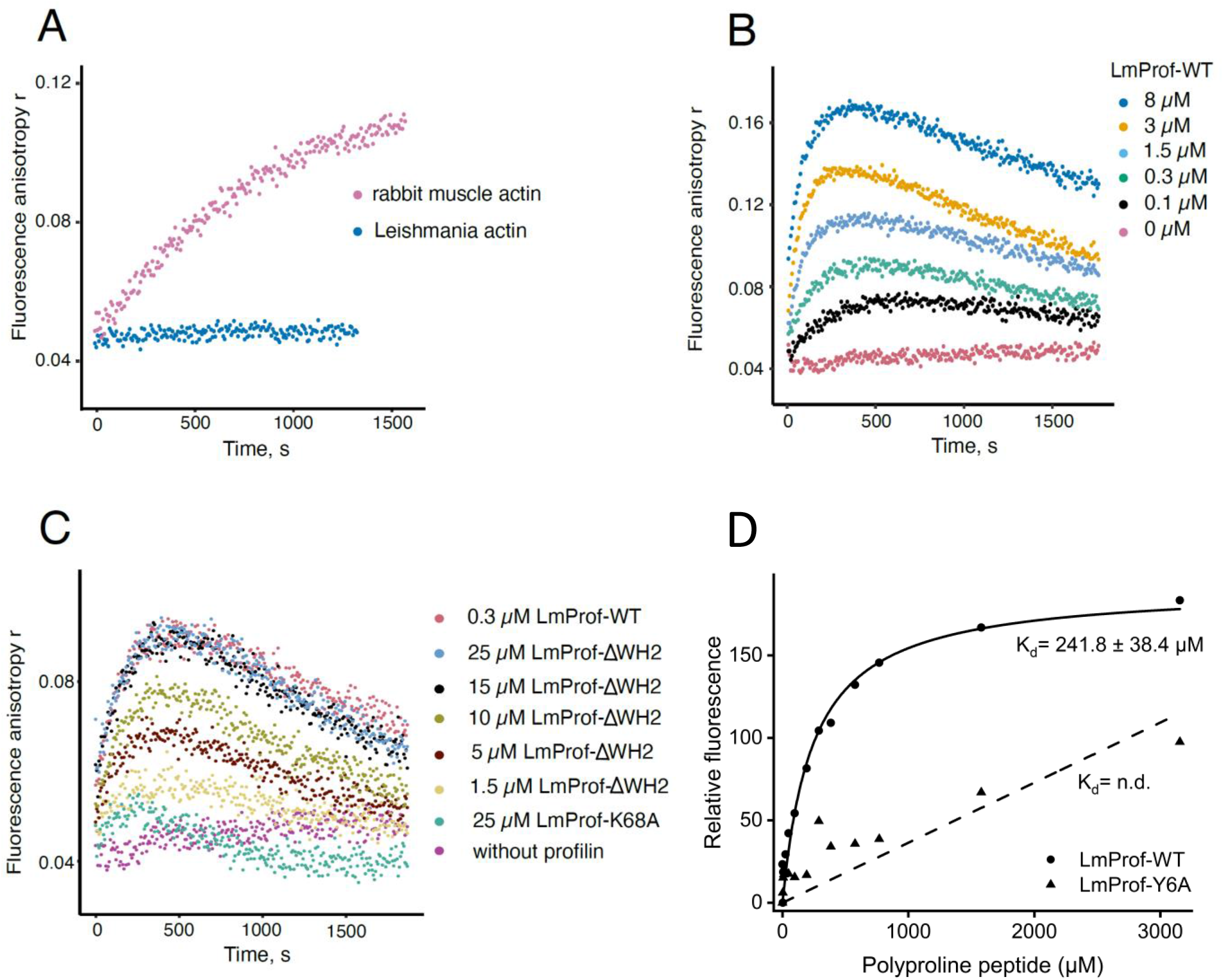
*Leishmania* profilin promotes nucleotide exchange on actin monomers, and binds poly-proline peptide through a conserved interface. **(A)** ATP-ATTO-488 fluorescence anisotropy experiment to compare nucleotide exchange kinetics of *Leishmania* and rabbit muscle monomeric actins. **(B)** ATP-ATTO-488 fluorescence anisotropy experiment to assess the effect of *Leishmania* profilin on nucleotide exchange kinetics of *Leishmania* monomeric actin. **(C)** ATP-ATTO-488 fluorescence anisotropy experiment to compare the activities of wild-type (WT) and two mutants of *Leishmania* profilin (ΔWH2 and K68A). **(D)** Tryptophan fluorescence assay to study the interaction of wild-type LmProfilin (LmProf-WT; solid circles) and LmProfilin-Y6A mutant (LmProf-Y6A; solid triangles) with a poly-L-proline peptide. Different concentrations of a decamer poly-L-proline peptide were mixed with 1 µM of profilins and the relative fluorescent intensity was measured. Data points are shown in symbols, the fitting curves in lines, and the obtained Kd value for wild-type profilin - poly-L-proline interaction is shown. The affinity of LmProfilin-Y6A to poly-L-proline was too low to be detected by this assay (n.d.= not determined).

Another important feature of profilins is their ability to bind to proline-rich proteins. Our structure shows that the poly-proline binding site is conserved in LmProfilin (Fig. S1B), and *L. donovani* profilin was recently reported to bind to poly-proline peptides in an affinity chromatography assay. However, the authors did not measure the binding affinity of the interaction, or map the residues critical for poly-proline binding [30]. Here, we used the change in intrinsic tryptophan fluorescence of profilin upon binding to poly-L-proline to determine the dissociation constant of LmProfilin from a poly-proline decamer. Based on this assay, wild-type LmProfilin binds the poly-L-proline peptide with an affinity (K_d_ of 241.8 ± 38.4 µM) that is similar to the ones reported for the interaction between poly-L-proline decamer and *Acanthamoeba* and human profilins [42]. Tyr6 in LmProfilin is located in the putative poly-proline binding site, and mutation of the corresponding tyrosine in *Schizosaccharomyces pombe* reduces affinity to poly-L-proline, and poorly complements the loss of profilin *in vivo* [38]. Similarly, replacing this tyrosine by alanine in LmProfilin (LmProfilin-Y6A) diminished binding to the poly-L-proline decamer to an undetectable level (Fig. 4D).

Collectively, these experiments demonstrate that LmProfilin binds poly-proline rich proteins and catalyzes the nucleotide exchange on actin monomers through interfaces that are conserved between human, yeast and parasite profilins. However, efficient nucleotide exchange also relies on the WH2-like motif, which increases the affinity of LmProfilin to actin monomers.

### Effects of *Leishmania* profilin on formin-catalyzed actin filament assembly

Profilins studied so far bind FH1 domains of formins, and can hence deliver actin monomers to formin FH2 domains to enhance actin filament nucleation and polymerization [24]. Due to the presence of the WH2-like motif in LmProfilin, we inspected whether *Leishmania* profilin could be docked to the barbed end of the FH2 domain-actin co-crystal structure from yeast. Interestingly, whereas mammalian profilins can be docked to the barbed end face of the terminal actin subunit of the FH2 domain-bound filament end [43] (Fig. 5A), the WH2-like motif of LmProfilin makes pronounced steric clashes with the FH2 domain of formin (Fig. 5B). This suggests that *Leishmania* profilin might not be able to work together with formin in promoting actin filament assembly. We thus examined the effects of LmProfilin on actin polymerization of LmActin by using pyrene-actin polymerization assay.

**Figure 5.**
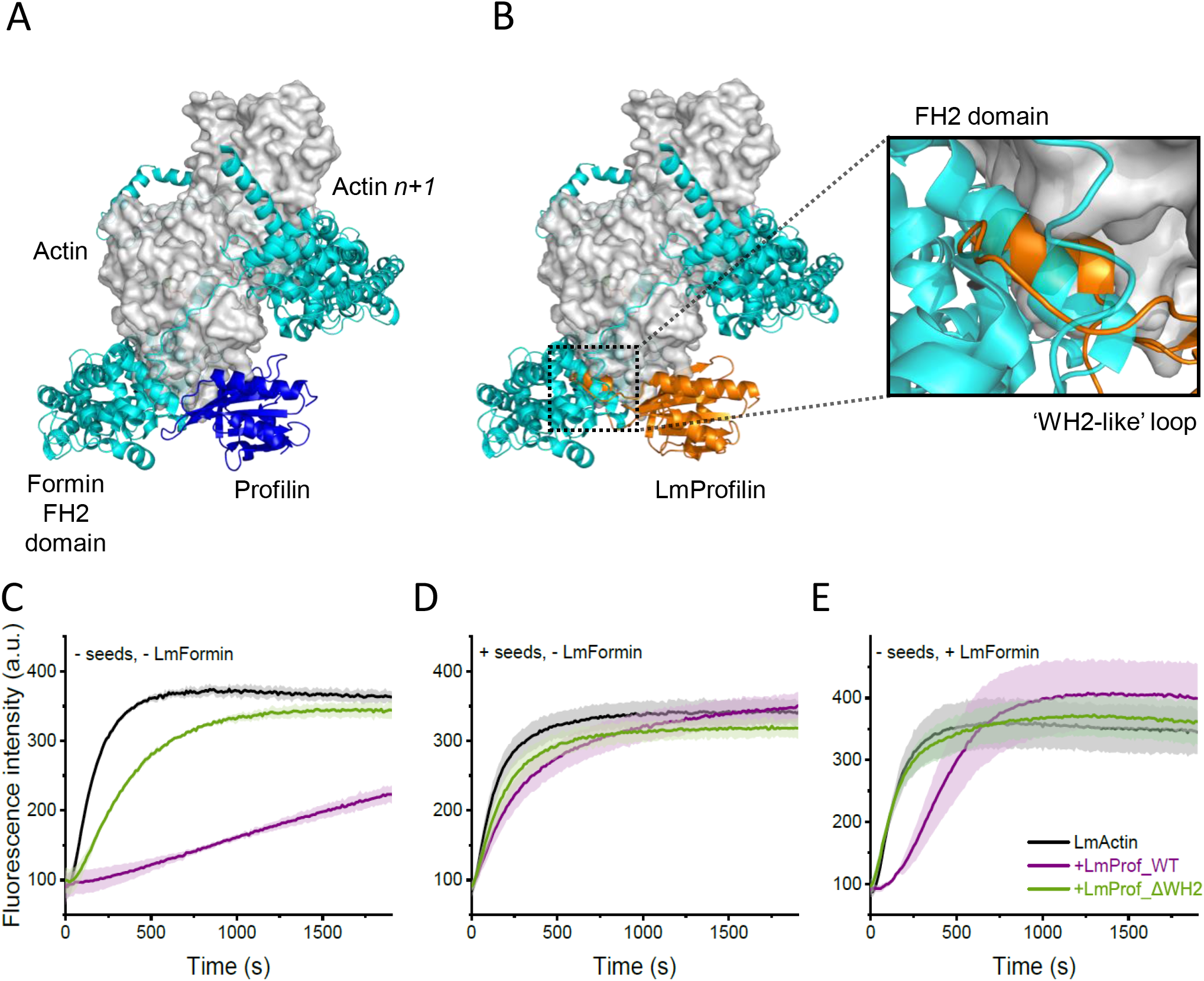
Effects of *Leishmania* profilin on formin-catalyzed actin filament assembly. **(A)** Human profilin-I (PDBID:2BTF) can be fitted to the barbed end of the ‘terminal’ actin subunit of the 2:2 formin FH2 domain:actin monomer structure (PDBID:1Y64) without steric clashes. **(B)** Docking the *Leishmania* profilin from our co-crystal structure to the FH2:actin structure results in major steric clashes between the profilin WH2 motif and the FH2 domain. **(C-E)** Pyrene-actin polymerization assays to monitor the effects of wild-type (LmProf_WT; purple curve), and LmProfilin-ΔWH2 (LmProf_ΔWH2; green curve) on spontaneous actin filament assembly (panel C), on actin filament assembly from actin-phalloidin seeds (panel D), and on actin filament assembly induced by LmFormin FH1-FH2 fragment (panel E). Final concentrations of actin (95% LmActin, 5% rabbit-pyrene actin) and profilin were 3 µM, and the concentrations of LmFormin FH1-FH2 and phalloidin seeds were 0.05 µM and 0.03 µM, respectively. Each curve depicts the average of four independent experiments with standard deviation shown in lighter color.

Because an earlier study on *Leishmania* actin dynamics demonstrated that rabbit muscle actin can co-polymerize with *Leishmania* actin [18], we performed pyrene-actin polymerization experiments by using a mix of 95 % LmActin and 5 % pyrene-labeled rabbit muscle actin. As demonstrated before [18], purified LmActin polymerized readily in the absence of any actin filament nucleators, and this is most likely due to rapid spontaneous nucleation of this actin (Fig. 5C). Addition of wild-type LmProfilin inhibited spontaneous assembly of actin filaments, similar to other profilins. Also, the profilin mutant lacking the WH2-like motif inhibited spontaneous actin filament assembly, although to a lesser extent, most likely due to the lower binding affinity of this mutant profilin to actin (Fig. 3; Fig. 5C). When the polymerization experiments were carried out in the presence of phalloidin-stabilized rabbit-actin seeds, the inhibition of actin assembly by profilin was mostly relieved, suggesting that LmProfilin does not significantly affect the incorporation of actin monomers to the pre-existing actin filament barbed ends (Fig. 5D). Interestingly, when the polymerization assay was carried in the presence of FH1-FH2 fragment of *L. major* formin-B (LmFormin), wild-type *Leishmania* profilin slowed down actin filament assembly by inducing an initial lag-phase to filament assembly, as well as by apparently reducing the maximal rate of filament polymerization. In contrast, *Leishmania* profilin lacking the WH2-like motif had no detectable effect on actin filament assembly in the presence of *Leishmania* formin, even at high concentrations to compensate weaker affinity (Fig. 5E; Fig. S5). Together these data demonstrate that, similar to other profilins, LmProfilin inhibits spontaneous actin filament nucleation, and maintains actin filament polymerization at filament barbed end. However, unlike profilins characterized from other organisms, LmProfilin is not compatible in promoting actin filament assembly with formins due to the presence of WH2-like motif, which makes a steric clash with formin at filament barbed end. Notably, the WH2-like motif is conserved in all trypanosomatid profilins suggesting a similar mode of action for profilin with formins in other trypanosomatid parasites.

### Interactions with actin and proline-rich proteins are important for the *in vivo* function of *Leishmania* profilin

To investigate the effects of the profilin mutations described above on *Leishmania*, we generated *L. mexicana* parasites expressing a range of profilin mutants (Fig S6, S7A). These include heterozygous (-/+) and homozygous (-/-) profilin knockouts, as well as *L. mexicana* knockin strains expressing mutant versions of profilin. In the knockin strains, the remaining *profilin* allele of the heterozygous (-/+) strain was replaced by Myc-tagged wild-type or mutant LmProfilin. Based on Western blot using polyclonal anti-LmProfilin antibody, the expression level of wild-type Myc-LmProfilin in the knockin strain (*profilin* -/WT) was slightly reduced as compared to the level of LmProfilin in the heterozygous (*profilin* -/+) strain (Fig. S7B). Because the polyclonal anti-LmProfilin antibody is likely to detect different mutant versions of LmProfilin with variable efficiency, we also probed the blot with anti-Myc antibody to compare the expression levels of wild-type and mutant profilins in the knockin strains. This demonstrated that all mutant Myc-LmProfilins were expressed either at similar or slightly higher protein levels as compared to the wild-type Myc-LmProfilin (Fig. S7B).

No effect on cell growth was observed in any profilin mutants, except a slight reduction in growth of the profilin null mutant (Fig. S8). Next, the effect of the profilin mutants on the uptake and trafficking of FM4-64, a lipophilic fluorescent dye, was examined. The flagellar pocket (FP) is the only known site for exocytosis and endocytosis in *Leishmania*, with FM4-64 initially accumulating at this point [44]. FM4-64 is then trafficked to the endocytic system and finally to the tubular lysosome, which runs along the anterior-posterior axis of the cell (Fig. 6A). We assessed the extent of uptake of FM4-64 for each profilin mutant and found a slower progression of FM4-64 trafficking in all profilin mutants compared to the wild-type profilin strain (*profilin* -/WT; Fig. 6B). The reduction of FM4-64 trafficking was slight in the proline-binding mutant (*profilin* -/Y6A), but more severe in the WH2 motif deletion (*profilin* -/ΔWH2) and the LL-SS (*profilin* -/LL-SS) mutants. There was also a significant reduction in FM4-64 trafficking in the actin binding interface mutants, especially the K86E mutant (*profilin* -/K86E). Together, these experiments demonstrate that in *L. mexicana*, profilin is not essential for viability in laboratory conditions, but is important for efficient endocytic trafficking. The rescue experiments also revealed that profilin’s ability to bind actin monomers and poly-proline proteins, as well as the presence of functional WH2-like motif contribute to its role in endosomal trafficking.

**Figure 6.**
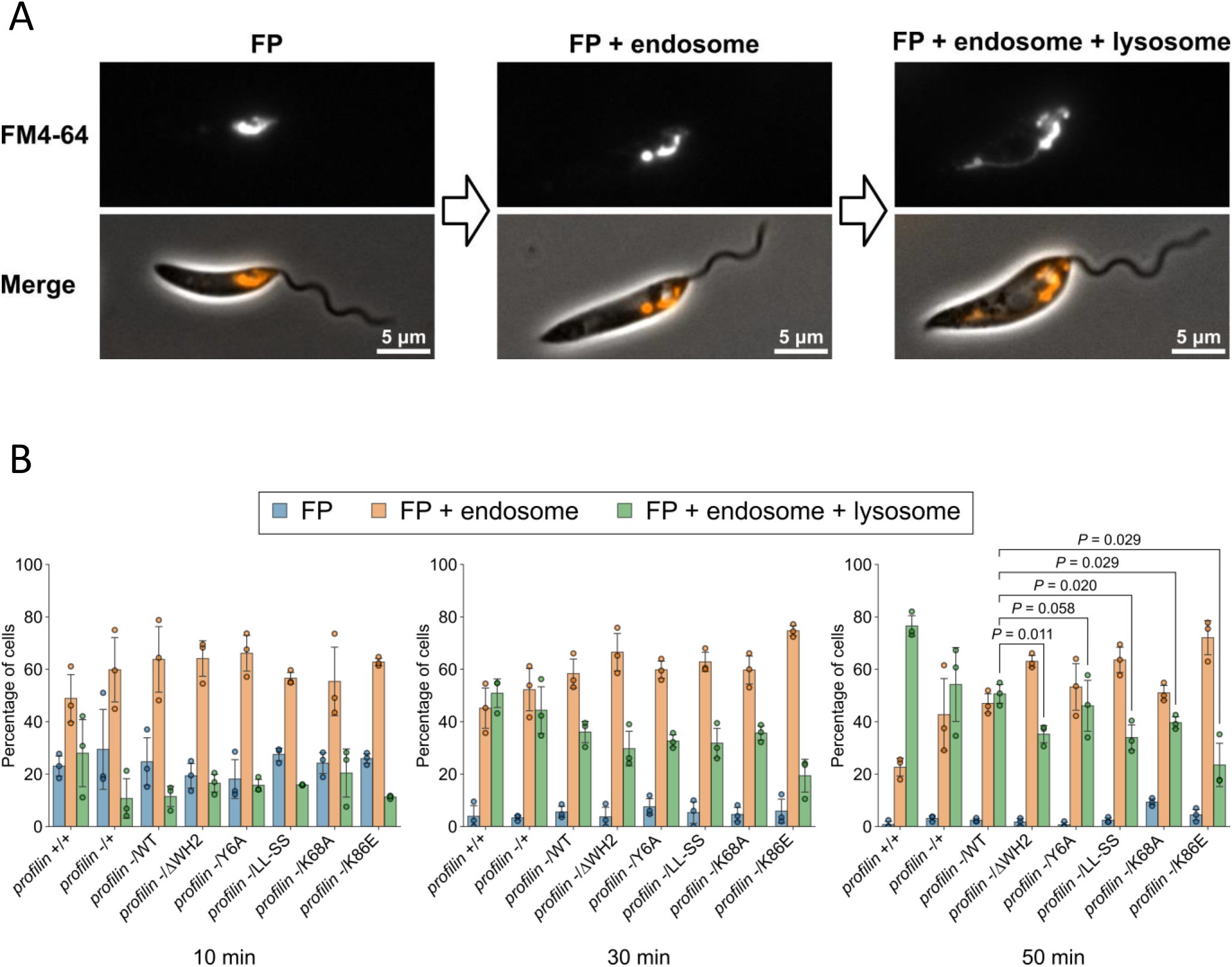
*Leishmania mexicana* parasites expressing profilin mutants have reduced FM4-64 trafficking. **(A)** Fluorescence and phase contrast microscopy images showing three major categories of FM4-64 localization after uptake into *L. mexicana* cells. Left: Flagellar pocket (FP). Middle: FP and endosome. Right: FP, endosome and lysosome. **(B)** FM4-64 endocytosis assay with profilin wild-type and mutant *L. mexicana* strains. Cells were chilled on ice for 20 min and then pulsed with FM4-64 for 1 min before imaging at 10, 30, 50 min time points. The number of cells in each of the three major categories of FM4-64 localization was counted at each time point. Data represent mean ± SD (n = 3 independent experiments). The dots represent individual measurement from three-independent experiments (33-62 cells were counted in each measurement). P-values calculated using two-tailed 787 Welch’s t-test.

## DISCUSSION

By determining the co-crystal structure of *L. major* profilin-actin complex, combined with biochemical work and studies on live parasites, our work uncovers the molecular principles by which actin dynamics are regulated by profilin in trypanosomatid parasites. This study also reveals important differences in the mechanisms by which trypanosomatid profilins associate with actin, as compared to actin-profilin interactions of other organisms studied so far.

Our structural and biochemical results provide evidence that the basic principles by which profilins interact with actin monomers are conserved across the eukaryotic domain. Similarly to eukaryotic profilins characterized so far, LmProfilin binds to the barbed end surface of actin, interacts with poly-proline motifs, and inhibits spontaneous actin filament nucleation (Figs. 1, 4, 5). Moreover, the ‘main’ actin-binding interface of profilin is conserved in evolution from mammals to trypanosomatid parasites. The principles of actin monomer binding are also conserved in even more distant profilins from Asgard archaea, whereas the mechanism by which archaea profilins associate with poly-proline appears divergent from eukaryotic profilin [36, 45]. However, in contrast to other profilins characterized so far, LmProfilin also harbors a WH2 domain–like insertion, which contributes to actin binding. In this context, it is important to note that also *Plasmodium* profilin harbors a structural insertion composed of two β-strands [35, 46, 47]. This insertion is important for malaria parasite motility, but it is located in a topologically different position of the protein and, based on MD simulation experiments, interacts with a different interface of actin as compared to the α-helical WH2-like motif of *Leishmania* profilin. Our structural and biochemical work provided evidence that the WH2-like motif of *Leishmania* profilin increases its affinity for actin monomers and thus helps to catalyze nucleotide exchange on actin monomers. On the other hand, the WH2-like motif makes a steric clash with formin FH2 domain at actin filament barbed end, and thus *Leishmania* profilin inhibits formin-mediated actin filament assembly, at least *in vitro*. In this context, it is important to note that in other organisms many activators of the Arp2/3 complex harbor WH2 domains [37], and it is possible that trypanosomatid profilins predominantly work together with the Arp2/3 complex to promote actin filament assembly.

The hypothesis on the role of trypanosomatid profilins in promoting Arp2/3-mediated actin filament assembly is also consistent with the phenotypes of *Leishmania* profilin knockout and knockin studies. Previous study on *L. donovani* heterozygous profilin mutant [30], and our present work on *L. mexicana* parasites expressing profilin mutants, demonstrated that profilin in trypanosomatid parasites is important for endocytic trafficking. Both endocytic internalization, as well as endosomal sorting are processes that rely on the Arp2/3 complex nucleated, branched actin filament networks [48]. Activation of the Arp2/3 complex during endocytic internalization is mainly catalyzed by WASP family proteins [49], whereas the Arp2/3 activation in endocytic sorting is catalyzed by the WASH protein complex [50, 51]. *Leishmania* genomes do not harbor clear homologues of WASP, but these organisms appear to express a protein, which displays weak sequence homology to the Arp2/3 interacting WASH-1 subunit of the WASH complex. Consistently, depletion of profilin results in defects in endosomal sorting, rather than endocytic internalization in *Leishmania* species. Our knockin studies also provided evidence that interactions with actin and poly-proline proteins, as well as the presence of a functional WH2-like motif are important for the function of *Leishmania* profilin in endocytic sorting. Thus, in addition to actin, profilin must also be able to interact with proline-rich proteins during endocytosis. It is important to note that endocytic Arp2/3 activators, including the subunits of the WASH complex, contain proline-rich segments, which are likely to bind profilin [52]. Thus, we speculate that endocytic sorting in *Leishmania* parasites relies on an Arp2/3-nucleated actin filament network, and that profilin delivers actin monomers to these filaments.

There are, however, important open questions about regulation of actin filament assembly in trypanosomatid parasites. In addition to the Arp2/3 complex, these parasites express formins. In animals and fission yeast, Arp2/3 and formins were shown to compete for a limited pool of actin monomers and profilin having an important role in controlling the balance between Arp2/3- and formin-catalyzed actin filament assembly [53, 54]. Therefore, it is likely that also trypanosomatid parasites have different actin filament populations. While the Arp2/3-nucleated actin filament structures in these organisms may contribute to endocytosis like they do in yeast and animal cells [48], formin-nucleated linear actin filament structures may contribute to some other cellular processes [55]. Thus, in the future it will be important to study the roles of the Arp2/3 complex and formins during different developmental stages of *Leishmania* parasites. Moreover, because *Leishmania* formins harbor proline-rich FH1 domains, it is likely that at least under certain circumstances *Leishmania* proflin can also deliver actin monomers to formins. It is, therefore, possible that the ‘formin-inhibitory effect’ of *Leishmania* profilin can be controlled by interactions with other proteins, or through specific post-translation modifications of profilin. Finally, it is interesting to note that both *Leishmania* actin, and especially actin-binding proteins display notable differences, both in their biochemistry and structures, compared to human actin and actin-binding proteins. Because actin is essential for viability of trypanosomatids [12], these differences, including the peculiar structural mechanism of *Leishmania* profilin-actin interaction identified here, could be applied for designing specific inhibitors against pathogenic trypanosomatid parasites.

## MATERIALS AND METHODS

### Cloning

The gene sequence of wild type *L. major* profilin (LmjF.32.0520) was taken from TriTrypDB database, codon optimized for bacterial expression, synthetized by TWIST Bioscience, and cloned into 3C/Precision cleavable double-tagged (Hisx6-GST) plasmid pCoofy3, a gift from Sabine Suppmann (Addgene plasmid # 43983 ; http://n2t.net/addgene:43983 ; RRID:Addgene_43983) [18, 56]. The mutants were prepared by site directed mutagenesis (see Tables S2 and S3 for details) following the QuikChange site-directed protocol (Agilent). For generation of knockin cell lines the pPLOT blast-mNG-blast plasmid was generated with recoded wild type and mutant profilin genes (synthetized by TWIST Bioscience) between the HindIII and SacI sites. A gene fragment encoding LmForminB (aa 536-1193) was codon optimized, synthesized by Integrated DNA Technologies (IDT) and cloned into the same pCoofy3 plasmid.

### Protein purification

*L. major* actin (LmActin; TriTrypDB ID: LmjF.04.1230) fused with human-β-thymosin and a His10x-tag at the C-terminus of actin was expressed in ExpiSf9 insect cells using the baculovirus system and subsequently purified by Ni-NTA affinity chromatography followed by gel filtration, as previously described [18]. Pure protein was flash-frozen in liquid nitrogen for long term storage or immediately further processed. β-thymosin-His10x tag was removed by cleavage with α-chymotrypsin. After quenching of cleavage reaction with PMSF, polymerization of LmActin was induced by adding EGTA and MgCl_2_, and filamentous actin was pelleted at 124,759 × g at 10°C for 1 h. Actin pellets were washed and resuspended in G-buffer (2 mM Tris pH 7.5, 0.5 mM β-mercaptoethanol, 0.2 mM CaCl_2_, 0.2 mM ATP) to a final concentration of ∼0.8 mg/ml, and dialyzed against G-buffer. Before assays, LmActin was ultracentrifuged for 1 h at 124,759 × g at 4°C and the upper two-thirds were collected to ensure only the presence of monomeric LmActin [18]. The final LmActin concentration ranges between 14-19 µM. *L. major* profilin (LmProfilin; TriTrypDB ID: LmjF.32.0520) wild-type and mutants were expressed as fusion proteins with an N-terminal double tag (His6x-GST). The recombinant proteins were purified using an approach reported before [18]. Briefly, *E*.*coli* BL21(DE3) (Merck Millipore) cells were grown at 22°C in LB autoinduction media (AIMLB0210, Formedium) supplemented with kanamycin (20 µg/mL) for ∼24 h. After lysis, recombinant proteins were first purified using a Ni-NTA column (GE HealthCare) and the His-GST tag was removed by cleavage with 3C-PreScision protease and subsequent incubation with Ni^2+^ beads to remove the uncleaved protein and the tag from the solution. The recombinant profilins were further purified by gel filtration, concentrated using Amicon Ultracentrifugal filters with MWCO of 3 kDa (Merck), aliquoted, flash-frozen in liquid nitrogen and stored at -75°C until use. For *L. major* formin-B (LmFormin; TriTrypDB ID: LmjF.24.1110), the construct consisted only of the FH1-FH2 domains and the C-tail (aa 536-1193) fused to a His6x-GST-tag at the N-terminus of the formin fragment. Expression and purification of the polypeptide was carried out in a similar way as described above and elsewhere [57], with slight modifications. Bacterial cells were harvested by centrifugation and lysed by sonication in the presence of lysozyme (0.5 mg/mL), DNase I (0.1 mg/mL) and protease inhibitors. The lysate was clarified by centrifugation and by passing through a 0.45 µm filter before loading it into a Ni-NTA column connected to an AKTA Pure instrument (GE Healthcare). After the protein binding to Ni-NTA beads, the column was washed with binding buffer (50 mM Tris-HCl, 300 mM NaCl, 10 mM imidazole, 3% glycerol, pH 7.5) and the protein was eluted with a linear gradient until reached 100% of elution buffer (50 mM Tris-HCl, 300 mM NaCl, 250 mM imidazole, 3% glycerol, pH 7.5). Peak fractions were pooled and concentrated by Amicon MWCO 50 kDa filters and loaded into a HiLoad 16/600 Superdex 200 column equilibrated with 20 mM HEPES pH 8.0 buffer, 50 mM NaCl, 3% glycerol, for gel filtration chromatography. Fractions corresponding to the desired protein were pooled and cleaved with 3C PreScision protease for 1 h at 4°C with gentle rotation. Glutathione-sepharose 4 beads (GE Healthcare) were added to remove the cleaved tags and non-cleaved protein for another 2 h at 4°C in a column under gravity flow. Protein was aliquoted and flash frozen for storage at -75°C until further use. Yeast profilin (Pfy1p) and mouse profilin-1 (PROF1) were expressed in Rosetta2(DE3)pLysS cells as fusion proteins with an N-terminal tag (6xHis-TEV). The recombinant proteins were batch purified on Nickel-Sepharose beads 6 Fast Flow (GE Healthcare) and eluted with 6xHis-TEV protease. Proteins were concentrated by Amicon filters MWCO 10 kDa, dialyzed for 2h at 4°C against storage buffer (20 mM Hepes pH 7.5, 50 mM KCl, 6% glycerol) and flash frozen for storage. Rabbit muscle actin was purified as described in [58].

### Crystallization and structure determination

LmProfilin and LmActin were mixed at at ∼ 1:1 molar ratio and 4.5 – 6.5 mg/ml concentration for sitting drop crystallization at Crystallization core facility (Institute of Biotechnology, HiLife). Hits were obtained from 0.1 M Bis-Tris pH 5.5, PEG4000 26% (w/v) and 0.2 M NaCl. Crystals were fished and cryo-protected with 15-20% glycerol for shipment to remote data collection at Diamond Light Source (beamline I03, Oxfordshire, Didcot, UK). The data were collected with 0.1° oscillation per frame and 0.010 s exposure time at a wavelength of 0.9762 Å and processed with autoPROC package, which utilizes XDS and AIMLESS for indexing, integration and scaling of the data [59-62]. Next, we used the sequences of *L. major* actin and *L. major* profilin as inputs for homologous model search molecular replacement with ARP/wARP classic model building web service [63, 64] (https://arpwarp.embl-hamburg.de/). After the initial solution, the model was finalized with manual curation in Coot [65] and rounds of refinement in Buster [66].

### Isothermal titration calorimetry (ITC)

Isothermal titration calorimetry (ITC) assays were performed in a Microcal-PEAQ instrument (Malvern Panalytical). Both LmActin and the proteins used for titration (wild-type LmProfilin and mutant versions) were dialyzed to G-buffer, and degassed under vacuum for 30 min before each experiment. Titrations were done at 22°C with LmActin (14-19 µM) loaded in the sample cell (250 µL) and injecting the ligand (150 µM in the syringe), with an initial 0.5 µL-injection followed by nineteen 2 µL-injections, each lasting 4 s and with 180 s of spacing between injections. The obtained thermograms were analyzed with the MicroCal PEAQ Analysis software using the single set of sites model to fit the curves. Heat due to dilution was corrected by control injections of LmProfilin into buffer.

### Tryptophan fluorescence assay

LmProfilins (wild type or mutant Y6A) at a final concentration of 1 µM were mixed at room temperature with different concentrations (at final concentrations from 3 to 3156 µM) of a decamer poly-L-peptide (CASLO ApS) in 20 mM Hepes pH 8.0, 50 mM NaCl buffer in a final volume of 110 µL. Change in intrinsic fluorescence intensity was measured in a Cary Eclipse Spectrophotometer (Agilent Technologies) at an excitation wavelength of 295 nm and range emission from 300-500 nm (5 nm-slit width). The normalized maximum intensity fluorescence (determined as the average of the five highest intensities) was plotted against the peptide concentration. To calculate the dissociation constant (K_d_), the data were analyzed in OriginPro and fit by nonlinear least-squares method using the OneSiteBind function.

### Pyrene-actin polymerization assays

Polymerization of LmActin was analyzed by the increase in fluorescence of pyrene-actin measured in a Cary Eclipse Fluorescence spectrophotometer, with an excitation wavelength of 365 nm and an emission wavelength of 407 nm. Before each experiment, LmActin was centrifuged at 124,759 × g for 60 min at 4 °C to remove possible oligomers. For each polymerization reaction, 60 µL of 1:10 final volume of 10x initiation buffer (20 mM Hepes, pH 7.4, 0.1 M KCl, 0.1 mM EGTA, 1 mM MgCl_2_, 0.2 mM ATP) containing either wild-type LmProfilin, or mutant LmProfilin ΔWH2 (in a final concentration of 3 µM), with or without LmFormin FH1-FH2 and phalloidin stabilized-actin seeds (at final concentrations of 0.05 µM and 0.03 µM, respectively) and G-buffer if needed, was combined with 60 *μ*l of monomeric LmActin (5% pyrene-rabbit actin) in G-buffer to yield a final concentration of 3 µM of actin in a final volume reaction of 120 µL. For the titration assays, the same conditions and concentrations for LmActin and LmFormin were used in combination with different concentrations (1, 3, 5, 10 µM) of wild-type LmProfilin or LmProfilin ΔWH2.

### Nucleotide exchange assay

Rabbit muscle or *L. major* actins were incubated with a 1:10 volume of 10x exchange buffer (100 mM Tris pH 8.0, 2.5 mM EGTA, 1 mM MgCl_2_ and 0.2 mM ADP) for 5 minutes on ice, and then dialyzed against G-ADP buffer (5mM Tris pH 7.5, 0.1 mM MgCl_2_, 0.02 mM ADP and 0.5 mM DTT) for 1h at 4°C. All exchange experiments reported in this article were initiated by incubation of G-actin (1 µM) with N^6^-(6-Amino)hexyl-ATP-ATTO-488 (0.1 µM; Jena Bioscience, Germany, ref. NU-805-488) in G+ME buffer (5mM Tris pH 7.5, 0.1 mM CaCl_2_, 0.5 mM DTT, 1 mM EGTA and 1 mM MgCl_2_) at room temperature. Anisotropy values were recorded by excitation at 504 nm and emission at 521 nm on a Safas Xenius XC spectrofluorimeter (Safas Monaco), using a kinetic acquisition mode available on the version 7.8.13.0 of the SP2000 software. Data were plotted with RStudio.

### *Leishmania* cell culture and generation of profilin knock in/out mutants

*L. mexicana* promastigotes expressing Cas9 nuclease and T7 RNA polymerase were grown at 28°C in M199 medium with 10% foetal calf serum, 40 mM HEPES-NaOH (pH 7.4), 26 mM NaHCO3 and 5 µg/ml haemin [67]. Cells were maintained in logarithmic growth. Profilin knockout constructs and guide RNAs were generated as described [67]. LeishGEdit was used to design primers for use with the knockout plasmid pTNeo. The plasmids (pPL1795-1800) were used as templates to generate knockin constructs that were transfected in alongside the profilin 5’ guide RNA template (Figure S6). The knockout and knockin constructs were transfected into 1×10^7^ cells resuspended in transfection buffer (200 mM Na_2_HPO_4_, 70 mM NaH_2_PO_4_, 15 mM KCl, 150 mM HEPES (pH 7.3) and 1.5 mM CaCl_2_), using programme X-001 on a Amaxa Nucleofector IIb. After electroporation cells were transferred into 10 ml of M199 and incubated at 28°C. After ∼6 hours, transfected cells were selected with appropriate drug (blasticidin – 5 µg/ml, G418 – 20 µg/ml) for 5-10 days before sub-culturing of successful transformants.

To confirm the knockout/knockin of profilin in mutant cells, PCR was performed on gDNA extracted using DNeasy Blood & Tissue kit (Qiagen).

### Endocytosis assays

A total of 5 × 10^6^ cells of log-phase *L. mexicana* promastigotes were incubated in M199 medium on ice for 20 min before final concentration of 5 µg/ml FM4-64 (Invitrogen; from a 1000 µg/ml stock solution in DMSO) was added for 1 min at room temperature (RT). Cells were centrifuged at 800 × g for 3 min at RT, resuspended in 600 µl of prewarmed M199 at 28°C and then divided into three tubes of 200 µl each. Each tube was incubated at 28°C and at each time point (10, 30 and 50 min), one of the tubes was centrifuged at 800 × g for 1 min at RT to concentrate cells for imaging with a Zeiss ImagerZ2 microscope with a 63× NA 1.4 objective and Hamamatsu Flash 4 camera. Captured cells were categorised according to the FM4-64 localisation (Fig. 6A; flagellar pocket (FP); FP and endosome; and FP, endosome, and lysosome).

### Western blots

A total of 4 × 10^7^ cells of log-phase *L. mexicana* promastigotes were harvested by centrifugation (800 × g for 7 min at RT). The cells were washed with 5 ml of PBS (137 mM NaCl, 2.7 mM KCl, 10 mM Na_2_HPO_4_, 1.8 mM KH_2_PO_4_) and resuspended in 1 ml of ice cold PBS with cOmplete, EDTA-free protease inhibitor cocktail (Roche). The cells were pelleted by centrifugation (10,000 × g for 2 min at RT) and resuspended in 200 µl of Laemmli buffer (2% SDS, 10% Glycerol, 60 mM Tris-HCl, 50 mM DTT, pH6.8) with cOmplete EDTA-free protease inhibitor cocktail. Cell lysates (approximately 4 × 10^6^ cell equivalents, without heating) were loaded and subjected to electrophoresis on 15% SDS-polyacrylamide gels and transferred to a nitrocellulose membrane (GE Healthcare) in transfer buffer (25 mM Tris, 192 mM Glycine, 20% Methanol) without SDS. The membrane was blocked in 5% skim milk in Tris buffered saline (20 mM Tris, 150 mM NaCl, pH7.5) with 0.1% (w/v) Tween 20 (TBST) at RT for 1 h, then probed overnight at 4°C with 1:500 dilution of guinea pig anti-LmProfilin antiserum (raised against recombinant LmProfilin by Pineda Antikörper-Service, Berlin, Germany) or with mouse anti-myc-tag antibody (clone 9E10 hybridoma supernatant, grown in Sunter lab) in blocking buffer. After washing with TBST, membranes were incubated at RT for 1 h with 1:1000 dilution of horseradish-peroxidase conjugated rabbit anti-guinea pig IgG secondary antibody (Invitrogen), or with 1:2500 dilution of horseradish-peroxidase conjugated donkey anti-mouse IgG secondary antibody (Jackson ImmunoResearch) in blocking buffer, washed in TBST and incubated with WesternBright Quantum (advansta). The membrane was visualised by G:BOX Chemi XRQ instrument (Syngene).

## ACKNOWLEDGMENTS

This work was supported by Academy of Finland (302161) and Jane and Aatos Erkko Foundation (4708679) to P.L., by the Wellcome Trust (221944/Z/20/Z) and Leverhulme Trust to J.D.S., the Fondation pour la Recherche Médicale (grant number EQU202103012764) to A.M., by the Ligue Contre le Cancer (fellowship number IP/SC-17531) to J.C., and by a JSPS Overseas Research Fellowship to R.Y. The facilities and expertise of the HiLIFE Crystallization unit at the University of Helsinki, a member of FINStruct and Biocenter Finland are gratefully acknowledged for crystallization services, and the Diamond Light Source for beamtime (proposal MX9951-28) at I03 beamline. We also thank Mirva Tirkkonen for excellent technical assistance.

## REFERENCES

1. Mann S, Frasca K, Scherrer S, Henao-Martinez AF, Newman S, Ramanan P, et al. A Review of Leishmaniasis: Current Knowledge and Future Directions. Curr Trop Med Rep. 2021;8:121–32.

2. World Health Organization. Leishmaniasis 2022 [December 19, 2022]. Available from: https://www.who.int/news-room/fact-sheets/detail/leishmaniasis.

3. Kaufer A, Ellis J, Stark D, Barratt J. The evolution of trypanosomatid taxonomy. Parasit 549 Vectors. 2017;10:287.

4. Sunter J, Gull K. Shape, form, function and Leishmania pathogenicity: from textbook descriptions to biological understanding. Open Biol. 2017;7(9). doi: 10.1098/rsob.170165.

5. Zinoviev A, Shapira M. Evolutionary conservation and diversification of the translation initiation apparatus in trypanosomatids. Comp Funct Genomics. 2012;2012:813718.

6. Gull K. The cytoskeleton of trypanosomatid parasites. Annu Rev Microbiol. 1999;53:629–55.

7. Berriman M, Ghedin E, Hertz-Fowler C, Blandin G, Renauld H, Bartholomeu DC, et al. The genome of the African trypanosome Trypanosoma brucei. Science. 2005;309:416–22.

8. de Souza W, Meza I, Martinez-Palomo A, Sabanero M, Souto-Padron T, Meirelles MN. Trypanosoma cruzi: distribution of fluorescently labeled tubulin and actin in epimastigotes. J Parasitol. 559 1983;69:138–42.

9. Mortara RA. Studies on trypanosomatid actin. I. Immunochemical and biochemical identification. J Protozool. 1989;36:8–13.

10. Blanchoin L, Boujemaa-Paterski R, Sykes C, Plastino J. Actin dynamics, architecture, and mechanics in cell motility. Physiol Rev. 2014;94:235–63.

11. Lappalainen P, Kotila T, Jegou A, Romet-Lemonne G. Biochemical and mechanical regulation of actin dynamics. Nat Rev Mol Cell Biol. 2022;23:836–52.

12. Garcia-Salcedo JA, Perez-Morga D, Gijon P, Dilbeck V, Pays E, Nolan DP. A differential role for actin during the life cycle of Trypanosoma brucei. EMBO J. 2004;23:780–9.

13. Tammana TV, Sahasrabuddhe AA, Bajpai VK, Gupta CM. ADF/cofilin-driven actin dynamics in early events of Leishmania cell division. J Cell Sci. 2010;123:1894–901.

14. Gupta CM, Ambaru B, Bajaj R. Emerging Functions of Actins and Actin Binding Proteins in Trypanosomatids. Front Cell Dev Biol. 2020;8:587685.

15. Kapoor P, Kumar A, Naik R, Ganguli M, Siddiqi MI, Sahasrabuddhe AA, et al. Leishmania actin binds and nicks kDNA as well as inhibits decatenation activity of type II topoisomerase. Nucleic 574 Acids Res. 2010;38:3308–17.

16. Akil C, Kitaoku Y, Tran LT, Liebl D, Choe H, Muengsaen D, et al. Mythical origins of the actin cytoskeleton. Curr Opin Cell Biol. 2021;68:55–63.

17. Pollard TD. Actin and Actin-Binding Proteins. Cold Spring Harb Perspect Biol. 2016;8.

18. Kotila T, Wioland H, Selvaraj M, Kogan K, Antenucci L, Jegou A, et al. Structural basis of rapid actin dynamics in the evolutionarily divergent Leishmania parasite. Nat Commun. 2022;13:3442.

19. Vizcaino-Castillo A, Osorio-Mendez JF, Ambrosio JR, Hernandez R, Cevallos AM. The complexity and diversity of the actin cytoskeleton of trypanosomatids. Mol Biochem Parasitol. 2020;237:111278.

20. Goldschmidt-Clermont PJ, Machesky LM, Doberstein SK, Pollard TD. Mechanism of the interaction of human platelet profilin with actin. J Cell Biol. 1991;113:1081–9.

21. Eads JC, Mahoney NM, Vorobiev S, Bresnick AR, Wen KK, Rubenstein PA, et al. Structure determination and characterization of Saccharomyces cerevisiae profilin. Biochemistry. 1998;37:11171–81.

22. Wolven AK, Belmont LD, Mahoney NM, Almo SC, Drubin DG. In vivo importance of actin nucleotide exchange catalyzed by profilin. J Cell Biol. 2000;150:895–904.

23. Watanabe N, Madaule P, Reid T, Ishizaki T, Watanabe G, Kakizuka A, et al. p140mDia, a mammalian homolog of Drosophila diaphanous, is a target protein for Rho small GTPase and is a ligand for profilin. EMBO J. 1997;16:3044–56.

24. Chesarone MA, DuPage AG, Goode BL. Unleashing formins to remodel the actin and microtubule cytoskeletons. Nat Rev Mol Cell Biol. 2010;11:62–74.

25. Courtemanche N. Mechanisms of formin-mediated actin assembly and dynamics. Biophys Rev. 2018;10:1553–69.

26. Chalkia D, Nikolaidis N, Makalowski W, Klein J, Nei M. Origins and evolution of the formin multigene family that is involved in the formation of actin filaments. Mol Biol Evol. 2008;25:2717–33. Epub 20081006.

27. Funk J, Merino F, Venkova L, Heydenreich L, Kierfeld J, Vargas P, et al. Profilin and formin constitute a pacemaker system for robust actin filament growth. Elife. 2019;8.

28. Wilson W, Seebeck T. Identification of a profilin homologue in Trypanosoma brucei by complementation screening. Gene. 1997;187:201–9.

29. Osorio-Mendez JF, Vizcaino-Castillo A, Manning-Cela R, Hernandez R, Cevallos AM. Expression of profilin in Trypanosoma cruzi and identification of some of its ligands. Biochem Biophys Res Commun. 2016;480:709–14.

30. Ambaru B, Gopalsamy A, Tammana TVS, Subramanya HS, Gupta CM. Actin sequestering protein, profilin, regulates intracellular vesicle transport in Leishmania. Mol Biochem Parasitol. 2020;238:111280.

31. Ambaru B, Gangadharan GM, Subramanya HS, Gupta CM. Profilin is involved in G1 to S phase progression and mitotic spindle orientation during Leishmania donovani cell division cycle. PLoS One. 2022;17:e0265692.

32. Schutt CE, Myslik JC, Rozycki MD, Goonesekere NC, Lindberg U. The structure of crystalline profilin-beta-actin. Nature. 1993;365:810–6.

33. Fedorov AA, Pollard TD, Almo SC. Purification, characterization and crystallization of human platelet profilin expressed in Escherichia coli. J Mol Biol. 1994;241:480–2.

34. Ezezika OC, Younger NS, Lu J, Kaiser DA, Corbin ZA, Nolen BJ, et al. Incompatibility with formin Cdc12p prevents human profilin from substituting for fission yeast profilin: insights from crystal structures of fission yeast profilin. J Biol Chem. 2009;284:2088–97.

35. Moreau CA, Bhargav SP, Kumar H, Quadt KA, Piirainen H, Strauss L, et al. A unique profilin-actin interface is important for malaria parasite motility. PLoS Pathog. 2017;13:e1006412.

36. Akil C, Robinson RC. Genomes of Asgard archaea encode profilins that regulate actin. Nature. 2018;562:439–43.

37. Dominguez R. The WH2 Domain and Actin Nucleation: Necessary but Insufficient. Trends Biochem Sci. 2016;41:478–90.

38. Lu J, Pollard TD. Profilin binding to poly-L-proline and actin monomers along with ability to catalyze actin nucleotide exchange is required for viability of fission yeast. Mol Biol Cell. 2001;12:1161–75.

39. Colombo J, Antkowiak A, Kogan K, Kotila T, Elliott J, Guillotin A, et al. A functional family of fluorescent nucleotide analogues to investigate actin dynamics and energetics. Nat Commun. 2021;12:548.

40. De La Cruz EM, Pollard TD. Nucleotide-free actin: stabilization by sucrose and nucleotide binding kinetics. Biochemistry. 1995;34:5452–61.

41. Kasai M, Nakano E, Oosawa F. Polymerization of Actin Free from Nucleotides and Divalent Cations. Biochim Biophys Acta. 1965;94:494–503.

42. Petrella EC, Machesky LM, Kaiser DA, Pollard TD. Structural requirements and thermodynamics of the interaction of proline peptides with profilin. Biochemistry. 1996;35:16535–43.

43. Schutt CE, Karlen M, Karlsson R. A structural model of the profilin-formin pacemaker system 639 for actin filament elongation. Sci Rep. 2022;12:20515.

44. Sunter JD, Yanase R, Wang Z, Catta-Preta Cmc, Moreira-Leite F, Myskova J, et al. Leishmania flagellum attachment zone is critical for flagellar pocket shape, development in the sand fly, and pathogenicity in the host. Proc Natl Acad Sci U S A. 2019;116:6351–60.

45. Survery S, Hurtig F, Haq SR, Eriksson J, Guy L, Rosengren KJ, et al. Heimdallarchaea encodes profilin with eukaryotic-like actin regulation and polyproline binding. Commun Biol. 2021;4:1024. 645

46. Bhargav SP, Vahokoski J, Kallio JP, Torda AE, Kursula P, Kursula I. Two independently folding units of Plasmodium profilin suggest evolution via gene fusion. Cell Mol Life Sci. 2015;72:4193–203.

47. Moreau CA, Quadt KA, Piirainen H, Kumar H, Bhargav SP, Strauss L, et al. A function of profilin in force generation during malaria parasite motility that is independent of actin binding. J Cell 650 Sci. 2020;134(5).

48. Kaksonen M, Roux A. Mechanisms of clathrin-mediated endocytosis. Nat Rev Mol Cell Biol. 2018;19:313–26.

49. Kaksonen M, Toret CP, Drubin DG. Harnessing actin dynamics for clathrin-mediated endocytosis. Nat Rev Mol Cell Biol. 2006;7:404–14.

50. Gomez TS, Billadeau DD. A FAM21-containing WASH complex regulates retromer-dependent sorting. Dev Cell. 2009;17:699–711.

51. Derivery E, Sousa C, Gautier JJ, Lombard B, Loew D, Gautreau A. The Arp2/3 activator WASH controls the fission of endosomes through a large multiprotein complex. Dev Cell. 2009;17:712–23.

52. Linardopoulou EV, Parghi SS, Friedman C, Osborn GE, Parkhurst SM, Trask BJ. Human subtelomeric WASH genes encode a new subclass of the WASP family. PLoS Genet. 2007;3(12):e237.

53. Rotty JD, Wu C, Haynes EM, Suarez C, Winkelman JD, Johnson HE, et al. Profilin-1 serves as a gatekeeper for actin assembly by Arp2/3-dependent and -independent pathways. Dev Cell. 2015;32:54–67.

54. Suarez C, Carroll RT, Burke TA, Christensen JR, Bestul AJ, Sees JA, et al. Profilin regulates F-actin network homeostasis by favoring formin over Arp2/3 complex. Dev Cell. 2015;32:43–53.

55. Michelot A, Drubin DG. Building distinct actin filament networks in a common cytoplasm. Curr Biol. 2011;21:R560–9.

56. Scholz J, Besir H, Strasser C, Suppmann S. A new method to customize protein expression vectors for fast, efficient and background free parallel cloning. BMC Biotechnol. 2013;13:12.

57. Kotila T, Wioland H, Enkavi G, Kogan K, Vattulainen I, Jegou A, et al. Mechanism of synergistic actin filament pointed end depolymerization by cyclase-associated protein and cofilin. Nat Commun. 2019;10:5320.

58. Spudich JA, Watt S. The regulation of rabbit skeletal muscle contraction. I. Biochemical studies of the interaction of the tropomyosin-troponin complex with actin and the proteolytic fragments of myosin. J Biol Chem. 1971;246:4866–71.

59. Vonrhein C, Flensburg C, Keller P, Sharff A, Smart O, Paciorek W, et al. Data processing and analysis with the autoPROC toolbox. Acta Crystallogr D Biol Crystallogr. 2011;67:293–302.

60. Kabsch W. Xds. Acta Crystallogr D Biol Crystallogr. 2010;66:125–32.

61. Evans PR, Murshudov GN. How good are my data and what is the resolution? Acta Crystallogr D Biol Crystallogr. 2013;69:1204–14.

62. Winn MD, Ballard CC, Cowtan KD, Dodson EJ, Emsley P, Evans PR, et al. Overview of the CCP4 suite and current developments. Acta Crystallogr D Biol Crystallogr. 2011;67:235–42.

63. Langer G, Cohen SX, Lamzin VS, Perrakis A. Automated macromolecular model building for X-ray crystallography using ARP/wARP version 7. Nat Protoc. 2008;3:1171–9.

64. Murshudov GN, Skubak P, Lebedev AA, Pannu NS, Steiner RA, Nicholls RA, et al. REFMAC5 for the refinement of macromolecular crystal structures. Acta Crystallogr D Biol Crystallogr. 2011;67:355–67.

65. Emsley P, Lohkamp B, Scott WG, Cowtan K. Features and development of Coot. Acta Crystallogr D Biol Crystallogr. 2010;66:486–501.

66. Blanc E, Roversi P, Vonrhein C, Flensburg C, Lea SM, Bricogne G. Refinement of severely incomplete structures with maximum likelihood in BUSTER-TNT. Acta Crystallogr D Biol Crystallogr. 2004;60:2210–21.

67. Beneke T, Madden R, Makin L, Valli J, Sunter J, Gluenz E. A CRISPR Cas9 high-throughput genome editing toolkit for kinetoplastids. R Soc Open Sci. 2017;4:170095.

68. Holm L. Dali server: structural unification of protein families. Nucleic Acids Res. 2022;50:W210–5.

69. Rebowski G, Boczkowska M, Drazic A, Ree R, Goris M, Arnesen T, et al. Mechanism of actin N-terminal acetylation. Sci Adv. 2020;6(15):eaay8793.

70. Kursula I, Kursula P, Ganter M, Panjikar S, Matuschewski K, Schuler H. Structural basis for parasite-specific functions of the divergent profilin of Plasmodium falciparum. Structure. 2008;16:1638–48.

71. Gaucher JF, Mauge C, Didry D, Guichard B, Renault L, Carlier MF. Interactions of isolated C-terminal fragments of neural Wiskott-Aldrich syndrome protein (N-WASP) with actin and Arp2/3 complex. J Biol Chem. 2012;287:34646–59.

72. Hertzog M, van Heijenoort C, Didry D, Gaudier M, Coutant J, Gigant B, et al. The beta-thymosin/WH2 domain; structural basis for the switch from inhibition to promotion of actin assembly. Cell. 2004;117:611–23.

73. Ferron F, Rebowski G, Lee SH, Dominguez R. Structural basis for the recruitment of profilin-actin complexes during filament elongation by Ena/VASP. EMBO J. 2007;26:4597–606.

74. Paavilainen VO, Oksanen E, Goldman A, Lappalainen P. Structure of the actin-depolymerizing factor homology domain in complex with actin. J Cell Biol. 2008;182:51–9.

75. Irobi E, Aguda AH, Larsson M, Guerin C, Yin HL, Burtnick LD, et al. Structural basis of actin sequestration by thymosin-beta4: implications for WH2 proteins. EMBO J. 2004;23:3599–608.

